# Exposure of Men to PTSD-Promoting Trauma Elevates Levels of Sperm miRNAs with Anxiety and Depression-Inducing Activities

**DOI:** 10.64898/2026.04.22.720211

**Authors:** Mitra Sadat Shirazi, Alexandre Champroux, Aidan Chen, Denny Sakkas, Tammy M. Scott, Emily Mellen, Abigail Kaija, Larisa Ryzhova, Lucy Liaw, Arturo Hernandez, Larry A. Feig

## Abstract

Chronically stressing male rodents can induce stress-specific epigenetic changes in sperm that contribute to altered offspring phenotypes. Whether similar phenomena occur in men is unclear. This study addresses this knowledge gap by analyzing sperm microRNAs (miRNAs) from 51 men exposed to various levels of adult trauma including crime, disaster, and physical or sexual violence, quantified by the Trauma History Questionnaire (THQ), a measure of risk for Post-Traumatic Stress Disorder (PTSD). Four sperm miRNAs, miR-532-3p, 491-5p, 375-3p and 361-3p correlated positively with men’s THQ scores, showing 4X to 130X over expression in sperm from the most highly traumatized men. These changes were independent of men’s adverse childhood experiences (ACEs), which we previously linked to decreased miR-34/449 in their sperm; and sperm miR-34/449 levels were not associated with THQ scores. Injecting these 4 miRNAs into fertilized mouse oocytes at levels comparable to those found in men reporting high THQ scores yielded offspring with elevated anxiety-and depression-like phenotypes. This finding differs from the stress related phenotypes we observed in offspring of mice fertilized by sperm with reduced levels of miR-34/449. Consistent with only a small subset of men with high THQ scores developing PTSD, we observed no statistically significant increase in overall anxiety or depression among this highly traumatized group, however there were indications of increased sleeplessness, appetite and concentration difficulties and negative self-concept among this group. Nevertheless, almost all men reporting high THQ scores had elevated levels of all 4 of these miRNAs in their sperm, suggesting these trauma-induced epigenetic changes may raise mental health risks in the offspring of men with only subtle mental health problems. Since ∼20 % of men report either THQ or ACE scores in the ranges linked here and in our earlier study to changes in sperm miRNAs that in mice lead to elevated levels of stress-related behaviors, a large human population with an elevated risk of transmitting stress-related traits to their offspring likely exists.

## Introduction

Evidence for epigenetic inheritance, where the effects of the environment are transmitted to offspring via epigenetic changes in their germ cells, is extensive in plants and worms, growing in rodents, and only suggestive in humans (for review see (1)). Much of the support for epigenetic inheritance in mammals derives from investigations into how exposing male mice to various types of chronic stress leads to stress-specific changes in sperm miRNA content that cause stress-specific phenotypes in their offspring (2). For example, our studies have shown that exposing adolescent mice to chronic social instability (CSI) stress, which involves introducing a foreign mouse to a cage twice a week for 7 weeks, leads to elevated anxiety and defective sociability specifically in female offspring (3). These transgenerational effects are due, at least in part, to the introduction into zygotes of suppressed sperm levels of members of the miR-34/449 family (4). In contrast, others have shown that exposing mice to chronic variable stress leads to elevated levels of 9 different sperm miRNAs that do not cause changes in these behaviors but instead blunt the HPA axis response to a stressor in both male and female offspring (5).

Obtaining comparable evidence in humans is difficult because of the limited ability to manipulate conditions. Nevertheless, evidence does exist. For example, exposure to famine can severely alter offspring health for multiple generations (6). Moreover, negative intergenerational effects stemming from events, such as the Holocaust (7), or a diagnosis such as PTSD have been observed (8, 9). Although epigenetic mechanisms have been proposed, no direct evidence has been presented, leaving these effects attributed mainly to parenting practices. However, altered upbringing cannot account for the observation that newborns of men who experienced early life trauma already display enhanced brain white matter (10). Twin studies revealed a substantial (∼50%) genetic component to the inherited risk from PTSD (8, 11). What is not sufficiently appreciated is that part of this genetic component may be epigenetic, because in these studies designed to identify “genetic” contributions by comparing twins reared apart, both offspring should be similarly affected by environmentally altered delivery of miRNAs from sperm of trauma-exposed fathers to zygotes.

In fact, changes in human sperm biochemistry have been observed to be associated with specific environmental stressors to which men are exposed (12). For example, the levels of 5 sperm miRNAs and 4 tRNAs were found to be responsive to the level of young men’s stressful experiences (13). We found suppressed levels of members of the miR-34/449 family in sperm of men exposed to abusive and/or dysfunctional family life as revealed by high scores on the Adverse Childhood Experience (**ACE**) survey (14). This finding has been subsequently confirmed for miR-34c by others using different surveys for early childhood trauma (15). Reductions in sperm miR-34c/449 levels in these men are comparable to what we observed in sperm of adolescent mice exposed to CSI stress, which we showed contributes to anxiety and defective sociability observed in their female offspring (16). Moreover, we found that CSI stress alters sperm miR-34/449 levels by suppressing the level of miR-34c in exosomes derived from a subset of brain astrocytes that normally travel through the blood to sustain miR-34/449 levels in sperm (17).

To directly test in the same pool of men for specificity in how sperm respond to various human traumas as it does in mice, we directly compared men who experienced adverse childhood experiences (ACEs) to men exposed to adult traumas such as crime, disaster, and physical or sexual violence, quantified using the standardized Trauma History Questionnaire (THQ) used to measure PTSD risk (18). Strikingly, we found a different set of sperm miRNA levels correlate with men reporting high THQ than those associated with men reporting high ACE scores, and these high THQ score associated sperm miRNA changes led to different stress related phenotypes in offspring of mice generated by injecting them at comparable levels into control fertilized zygotes. This adds support for similar mechanisms being involved in how stress-specific sperm miRNA changes contribute to distinct epigenetic inheritance in both mice and men.

## Results

We expanded our previous study on how sperm miRNA content is altered in men by the extent of their **A**dverse **C**hildhood **E**xperiences (**ACEs**) (19) by now also having them complete the **T**rauma **H**istory **Q**uestionnaire. The **THQ** (18) is a widely used instrument designed to assess an individual’s adult history of traumatic events, specifically those that could meet the A1 stressor criterion for PTSD. It includes 24 yes/no questions covering three main areas of trauma exposure: (a) victimization through crime-related events, (b) general disasters and traumatic experiences (e.g., injury, natural disasters, witnessing death), and (c) unwanted physical and sexual experiences. The questionnaire not only records the specific types of traumas experienced but also the frequency of each event. Previous studies have shown a correlation between higher THQ scores and increased risk of PTSD symptomatology, as well as depression, in both men and women (20, 21).,

The participants were on average 36.4 years of age, with a mean BMI of 29.6 kg/m2. The demographic, behavioral, and sperm characteristics of study participants are summarized in **Supplemental Table 1**, stratified by THQ score (low, medium, and high). There were no clinically significant differences noted across the groups defined by THQ score for most parameters. Sperm count and motility, were also well within normal limits according to these guidelines for all groups (count ≥15 mil/mL; motility ≥40%). Although a morphology score of 4% is still considered normal according to the most recent World Health Organization guidelines for semen analysis parameters, there were some volunteers in each group presenting a lower score.

As a primary screen to find potential candidate miRNAs whose expression in sperm correlates with THQ score, we chose sperm samples from five men with the highest (8–14) and randomly selected from five men with the lowest (0) scores (see **Table 1**) and performed micro-RNA sequencing on each sample (**Fig. 1**). The volcano plot shows miRNAs with significant differences in the high THQ group (**Fig 1A, B**). miR-361-3p, miR-491-5p, miR-532-3p, miR-99b-5p, miR-99a-5p, and miR-193b-3p, were up-regulated, while miR-7-5p, miR-147b-5p, miR-424-3p and miR-548av-5p were down-regulated.

**Figure 1:**
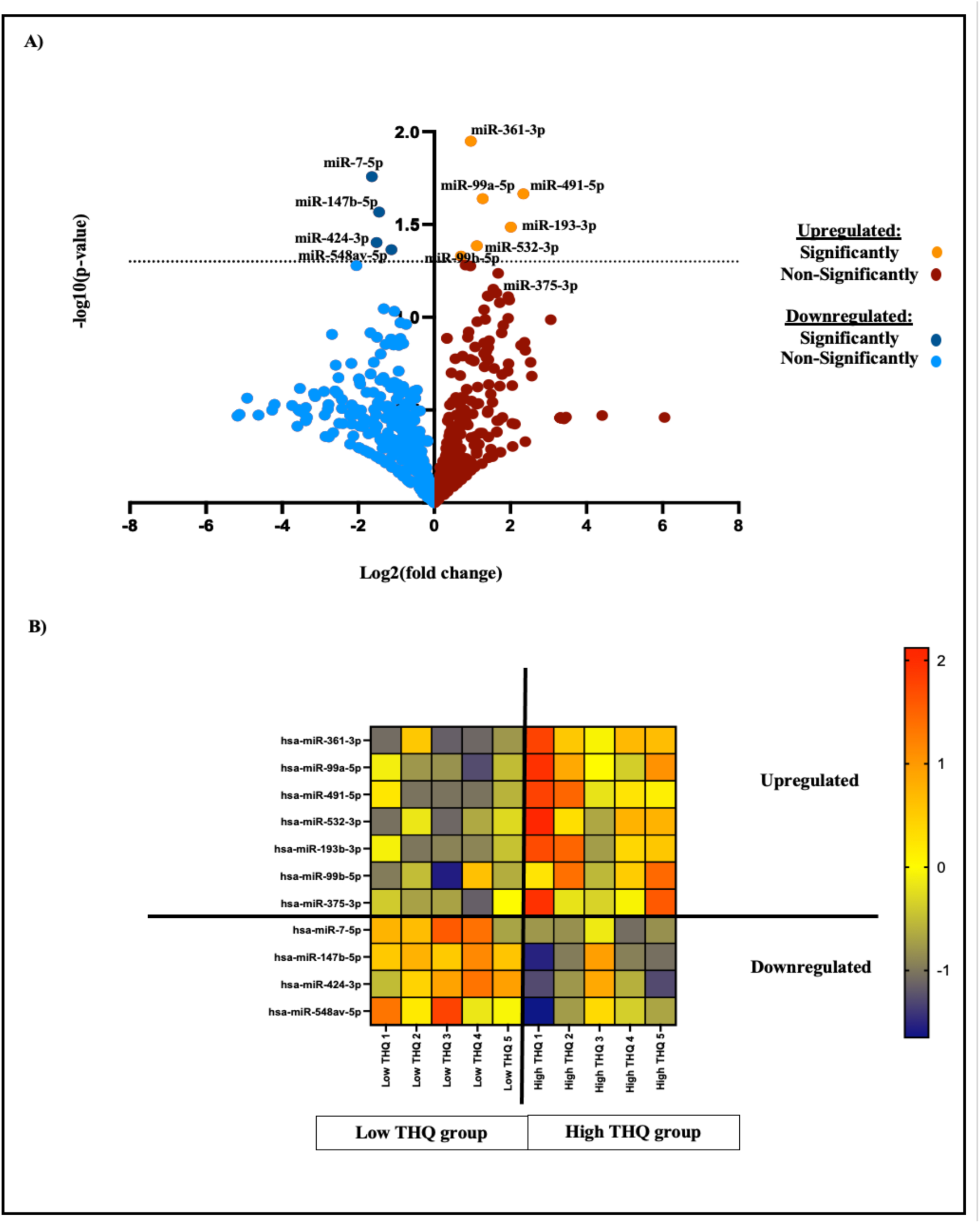
miRNA sequencing analysis between 5 low THQ and 5 high THQ. **A)** Volcano plot of differentially expressed human mature miRNA in 5 low THQ vs 5 high THQ sperm samples. The horizontal line () indicates the threshold of a 0.05 FDR value. The orange and the dark blue points lying in the top right and top left sectors are significantly upregulated and downregulated, respectively, in high THQ versus low THQ (FDR < 0.05). **B)** Heatmap of the 7 miRNAs upregulated significantly and 4 miRNAs downregulated between the 5 low and 5 high THQ samples. The red indicates higher miRNA expression level, and the blue shows the lowest miRNA expression level.

**Table 1:** Trauma History Questionnaire (or THQ). Compiled data for all the 51 human samples used in this study on the trauma history questionnaire (or THQ). All 24 questions with the number of n and the percentage for each group.

| Category | Variable n (%) | Whole samples (n=51) | Low THQ (0-2) N=19 | Medium THQ (3-7) N=25 | High THQ (≥8) N=7 |
| --- | --- | --- | --- | --- | --- |
| Crime-related events | Q1. Has anyone ever tried to take something directly from you by using force or the threat of force, such as a stick-up or mugging? | 7 (13.7) | 0 (0) | 6 (26) | 1 (14.3) |
|  | Q2. Has anyone ever attempted to rob you or actually robbed you (i.e., stolen your personal belongings)? | 14 (27.5) | 0 (0) | 9 (36) | 5 (71.5) |
|  | Q3. Has anyone ever attempted to or succeeded in breaking into your home when you were not there? | 7 (13.7) | 0 (0) | 7 (28) | 0 (0) |
|  | Q4. Has anyone ever attempted to or succeed in breaking into your home while you were there? | 6 (11.8) | 1 (5.3) | 3 (12) | 2 (28.6) |
| General disaster and trauma | Q5. Have you ever had a serious accident at work, in a car, or somewhere else? | 11 (21.6) | 3 (15.8) | 4 (16) | 4 (57.1) |
|  | Q6. Have you ever experienced a natural disaster such as a tornado, hurricane, flood or major earthquake, etc., where you felt you or your loved ones were in danger of death or injury? | 6 (11.8) | 0 (0) | 5 (20) | 1 (14.3) |
|  | Q7. Have you ever experienced a "man-made" disaster such as a train crash, building collapse, bank robbery fire, etc., where you felt you or your loved ones were in danger of death or injury? | 4 (7.8) | 0 (0) | 2 (8) | 2 (28.6) |
|  | Q8. Have you ever been exposed to dangerous chemicals or radioactivity that might threaten your health? | 4 (7.8) | 1 (5.3) | 2 (8) | 4 (7.8) |
|  | Q9. Have you ever been in any other situation in which you were seriously injured? | 8 (15.7) | 1 (5.3) | 4 (16) | 3 (42.9) |
|  | Q10. Have you ever been in any other situation in which you feared you might be killed or seriously injured? | 12 (23.5) | 1 (5.3) | 5 (20) | 6 (85.7) |
|  | Q11. Have you ever seen someone seriously injured or killed? | 12 (23.5) | 0 (0) | 7 (28) | 5 (71.4) |
|  | Q12. Have you ever seen dead bodies (other than at a funeral) or had to handle dead bodies for any reason? | 17 (33.3) | 1 (5.3) | 11 (44) | 5 (71.5) |
|  | Q13. Have you ever had a close friend or family member murdered, or killed by a drunk driver? | 3 (5.9) | 0 (0) | 3 (12) | 0 (0) |
|  | Q14. Have you ever had a spouse, romantic partner, or child die? | 1 (2) | 0 (0) | 1 (4) | 0 (0) |
|  | Q15. Have you ever had a serious or life-threatening illness? | 2 (3.9) | 0 (0) | 1 (4) | 1 (14.3) |
|  | Q16. Have you ever received news of a serious injury, life-threatening illness, or unexpected death of someone close to you? | 27 (52.9) | 5 (26.3) | 17 (68) | 5 (71.4) |
|  | Q17. Have you ever had to engage in combat while in military service in an official or unofficial war zone? | 3 (5.9) | 0 (0) | 1 (4) | 2 (29.8) |
| Physical and Sexual Experiences | Q18. Has anyone ever made you have intercourse or oral or anal sex against your will? | 0 (0) | 0 (0) | 0 (0) | 0 (0) |
|  | Q19. Has anyone ever touched private parts of your body, or made you touch theirs, under force or threat? | 0 (0) | 0 (0) | 0 (0) | 0 (0) |
|  | Q20. Other than incidents mentioned in questions 18 and 19, have there been any other situations in which another person tried to force you to have an unwanted sexual contact? | 2 (3.9) | 1 (5.3) | 1 (4) | 0 (0) |
|  | Q21. Has anyone, including family members or friends, ever attacked you with a gun, knife, or some other weapon? | 2 (3.9) | 0 (0) | 1 (4) | 1 (14.3) |
|  | Q22. Has anyone, including family members or friends, ever attacked you without a weapon and seriously injured you? | 3 (5.9) | 0 (0) | 0 (0) | 3 (42.9) |
|  | Q23. Has anyone in your family ever beaten, spanked, or pushed you hard enough to cause injury? | 0 (0) | 0 (0) | 0 (0) | 0 (0) |
|  | Q24. Have you experience any other extraordinarily stressful situation or event that is not covered above? | 5 (9.8) | 0 (0) | 3 (12) | 2 (28.6) |

But when we quantified each of these miRNAs in all 51 samples by qPCR and graphed them based on THQ score in **Fig. 2**, only three miRNAs, miR-532-3p, miR-491-5p and miR-375-3p, showed statistically significant positive correlations with all THQ scores. miR-361-3p fell just below standard significance (R=0.2578, p=0.06). (miR-99b-5p, miR-147b-5p, miR-424-3p and miR-548-5p were expressed at extremely low levels, making it difficult to test for a correlation).

**Figure 2:**
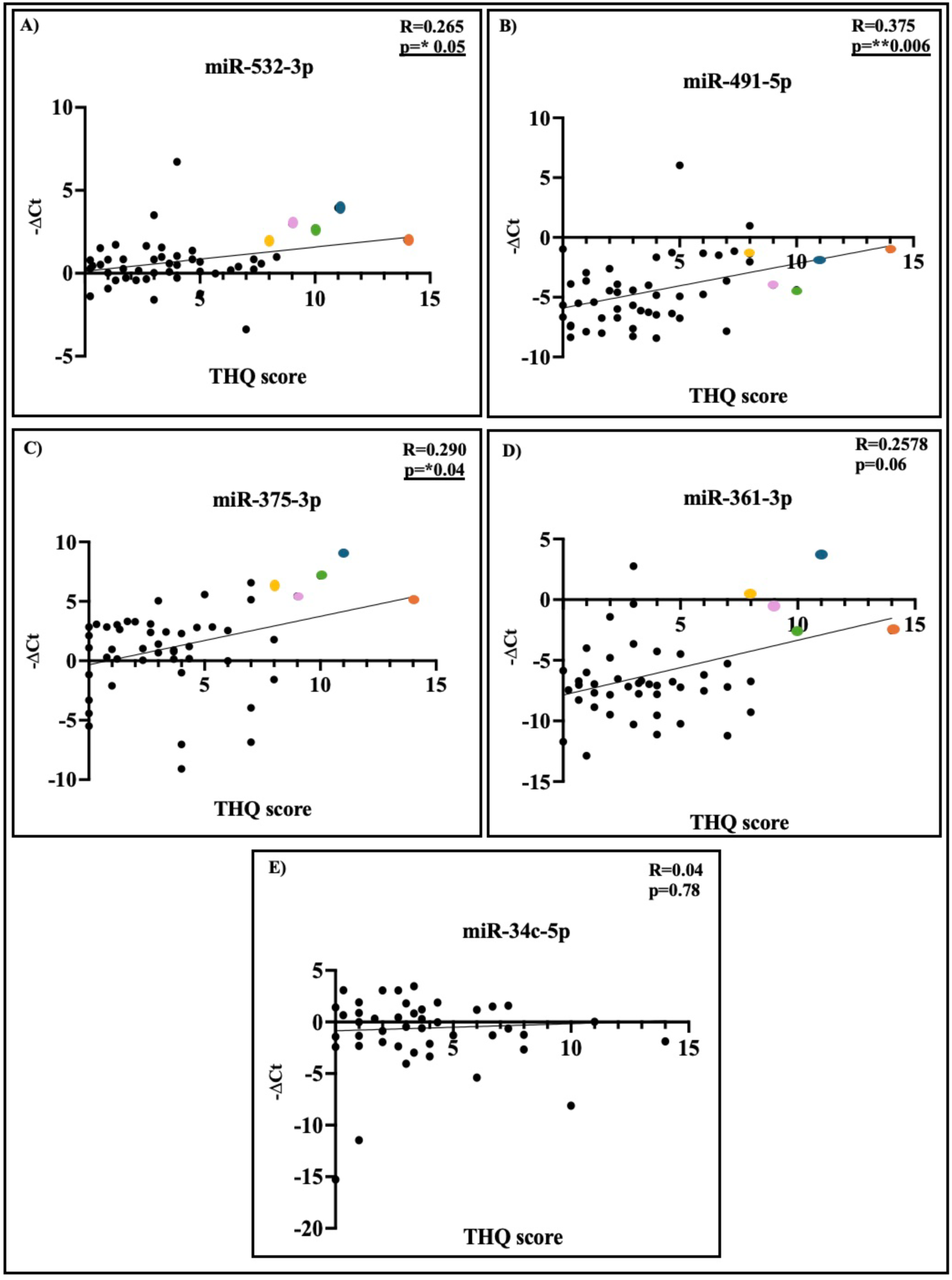
THQ score positively correlates with expression of miR-532-3p, miR-491-5p miR-375-3p and miR-361-3p, but not miR-34c-5p in human sperm samples. A-D) Analysis by qPCR of miR-532-3p, miR-491-5p, miR-375-3p and miR-361-3p in all human sperm samples, normalized to the -ΔCt for each individual point using miR-192-5p as the internal control for all samples. N=51, R= Spearman’s coefficient, with p values listed. Trend lines represent single-variable linear regression. Each color point represents results from the same volunteer. **E)** Same with miR-34c-5p. Each point represents an individual sample.

When we compared the levels of the 4 THQ-correlating miRNAs in sperm from men with the 5 highest with 5 representative men with THQ scores of zero, (**Fig. 3**) striking significant elevations in the levels of these miRNAs were revealed, with enhanced levels of 4X, 5X, 50X and 130X for miR-532-3p, miR-491-5p, miR-375-3p and miR-361-3p, respectively. Further enhancing the likelihood (i.e. non-randomness) these findings are significant is the fact that the 5 volunteers with the highest scores (each labeled with a different colored dot in **Fig. 2A-D)**) had elevated levels of all 4 of these miRNAs.

**Figure 3:**
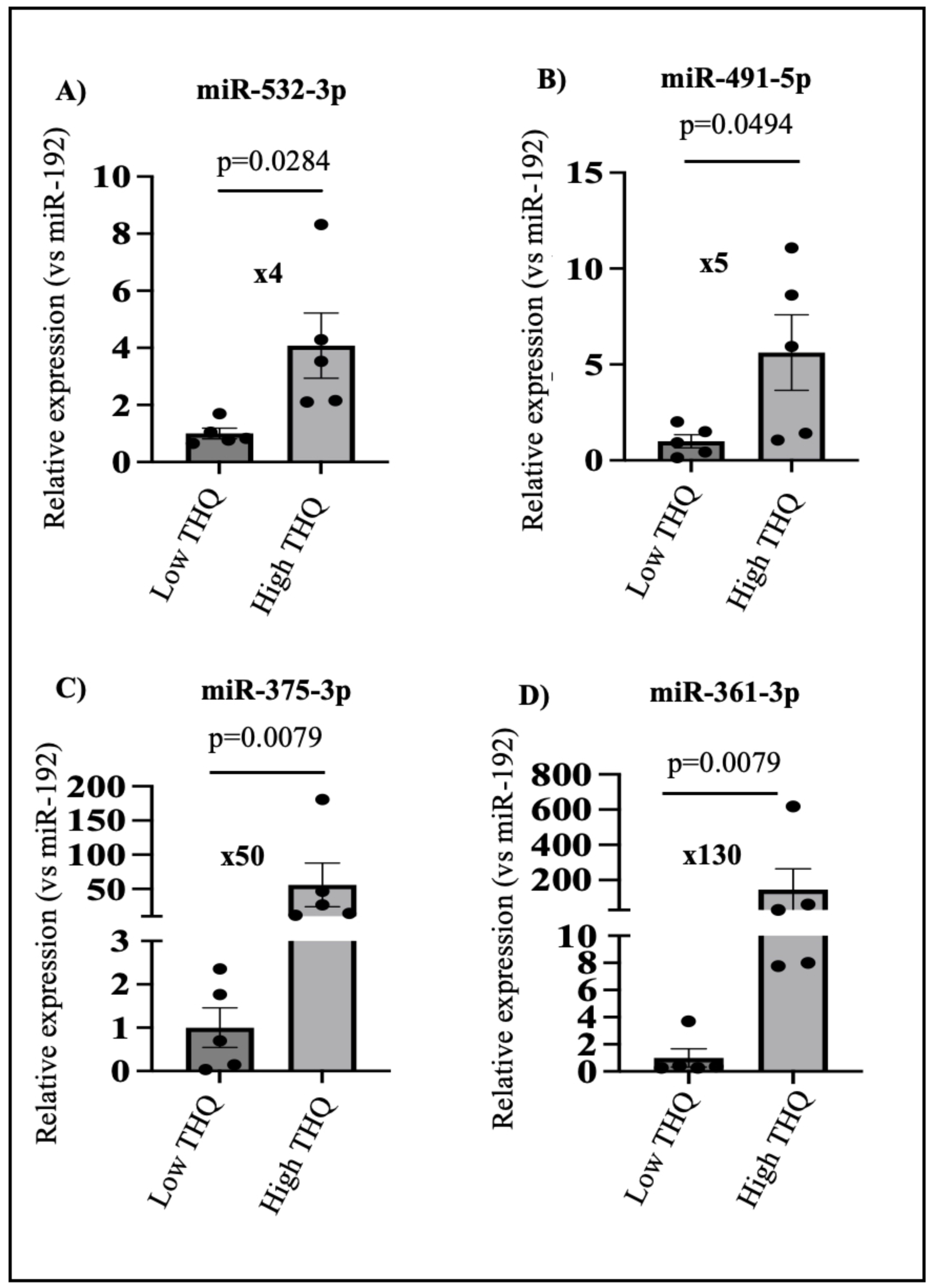
Increase of sperm miRNAs expression in men with extensive trauma history (or THQ). A-D) qPCR analysis of miR-532-3p, miR-491-5p, miR-375-3p and miR-361-3p in sperm RNA from low THQ group (score 0-2, n=5) and high THQ group (score ≥8, n=5). Data represented as mean ± SEM. All values are compared to the internal control miR-192-5p that does not change enough by any perturbation to significantly alter the conclusions and control ratios are set to 1. A and D, Unpaired t-test and B and C Mann-Whitney Test. The value X represents the mean difference between low THQ and high THQ.

We also compared the levels of these 4 miRNAs to these volunteers’ ACE scores, which in our previous study negatively correlated with sperm levels of miR-34/449, but found no correlation (**Table 2** and **Fig. 4A-D**). Moreover, levels of miR-34/449 did not correlate with THQ scores (**Fig. 2E**). These results are the first to directly compare two different trauma paradigms in the same set of men for their effects on sperm miRNA content to document that just as many have found in mice, sperm miRNA changes strongly depend upon the type of stress imposed.

**Figure 4:**
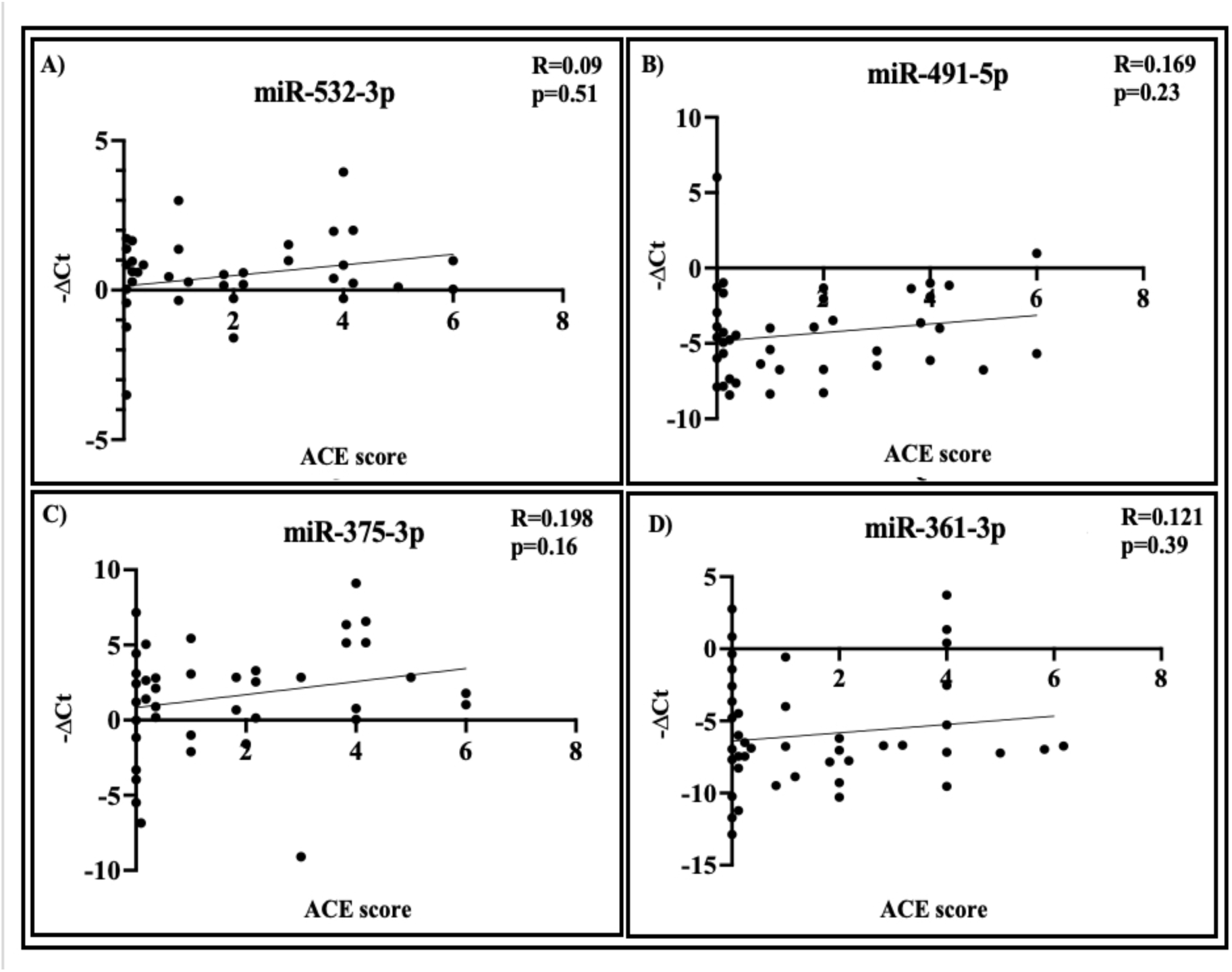
ACE score does not correlate with miRNAs expression in human sperm. **A-D**) miR-532-3p, miR-491-5p, miR-375-3p and miR-361-3p in all human sperm samples based on ACE score, normalized to the -ΔCt. Trend lines represent single-variable linear regression. N=51, R= Spearman’s coefficient, with p values listed. Each point represents an individual sample.

**Table 2:** Adverse Childhood Experiences (or ACE). Compiled data for all the 51 human samples used in this study on the Adverse Childhood Experiences (or ACE). All 10 questions with the number of n and the percentage for each group.

| Variable n (%) | Whole samples (n=51) | ACE 0 (n=28) | ACE 1 (n=5) | ACE 2 (n=6) | ACE 3 (n=2) | ACE 4 (n=7) | ACE 5 (n=1) | ACE 6 (n=2) |
| --- | --- | --- | --- | --- | --- | --- | --- | --- |
| Q1. Did a parent or other adult in the household often ... Swear at you, insult you, put you down, or humiliate you? Or Act in a way that made you afraid that might be physically hurt? | 9 (17.6) | 0 (0) | 1 (20) | 1 (16.7) | 0 (0) | 5 (71.4) | 1 (100) | 2 (100) |
| Q2. Did a parent or other adult in the household often ... Push, grab, slap, or throw something at you? Or ever hit you so hard that you had marks or were injured? | 4 (7.8) | 0 (0) | 0 (0) | 0 (0) | 1 (50) | 2 (28.6) | 0 (0) | 1 (50) |
| Q3. Did an adult or person at least 5 years older than you ever... touch or fondle you or have you touch their body in a sexual way? Or try to or actually have oral, anal, or vaginal sex with you? | 0 (0) | 0 (0) | 0 (0) | 0 (0) | 0 (0) | 0 (0) | 0 (0) | 0 (0) |
| Q4. Did you often feel that... no one in your family loved you or thought you were important or special? Or your family didn't look out for each other, feel close to each other, or support each other? | 3 (5.9) | 0 (0) | 0 (0) | 0 (0) | 0 (0) | 3 (42.9) | 0 (0) | 1 (50) |
| Q5. Did you often feel that ... You didn't have enough to eat, had to wear dirty clothes, and no one to protect you? Or your parents were too drunk or high to take care of you or take you to the doctor if you needed it? | 4 (7.8) | 0 (0) | 1 (20) | 2 (33.3) | 0 (0) | 0 (0) | 0 (0) | 1 (50) |
| Q6. Were your parents ever separated or divorced? | 16 (31.4) | 0 (0) | 1 (20) | 4 (66.7) | 2 (100) | 6 (85.7) | 1 (100) | 2 (100) |
| Q7. Was your mother or stepmother: often pushed, grabbed, slapped, or had something to throw at her? Or sometimes or often kicked, bitten, hit with a fist, or hit with something hard? Or ever repeatedly hit over at least a few minutes or threatened with a gun or knife? | 2 (3.9) | 0 (0) | 1 (20) | 0 (0) | 0 (0) | 1 (14.3) | 0 (0) | 1 (50) |
| Q8. Did you live with anyone who was a problem drinker or alcoholic or who used street drugs? | 14 (27.5) | 0 (0) | 1 (20) | 3 (50) | 2 (100) | 6 (85.7) | 1 (100) | 2 (100) |
| Q9. Was a household member depressed or mentally ill or did a household member attempt suicide? | 7 (13.7) | 0 (0) | 0 (0) | 2 (33.3) | 0 (0) | 3 (42.9) | 1 (100) | 1 (50) |
| Q10. Did a household member go to prison? | 5 (9.8) | 0 (0) | 0 (0) | 0 (0) | 1 (50) | 2 (28.6) | 1 (100) | 1 (50) |

Those reporting high THQ scores are predisposed to PTSD, so we included surveys of volunteers’ current feelings of anxiety and depression using the State-Trait Anxiety Inventory (22) and Patient Health Questionnaire (PHQ-8)(23), respectively (**Tables 3 and 4**). Consistent with previous findings showing only a small fraction of people scoring high on the THQ survey suffer from PTSD, statistically significant elevation in overall anxiety and depression was not found in the small group of 5-10 men reporting the highest THQ scores (**Fig. 5**). However, item-level descriptive statistics reveal higher mean scores on items related to difficulty sleeping (1.6 in the high THQ group vs. 0.9 and 0.8 in the low and medium groups, respectively), appetite changes (1.1 in high THQ vs. 0.65 low THQ and 0.60 medium THQ), negative self-concept (0.9 in high THQ vs. 0.55 in low THQ and 0.60 medium THQ), difficulty concentrating (1.3 in high THQ vs 0.6 low THQ and 0.8 medium THQ), and psychomotor agitation/retardation (0.7 in high THQ vs 0.1 low THQ and 0.2 medium THQ (see **Table 3** questions 3, 5, 6, 7, 8). These emotional and behavioral changes, though too subtle to distinguish high from medium/low THQ participants on a clinical screener, could explain the severe changes of expression displayed by their sperm years after experiencing trauma.

**Figure 5:**
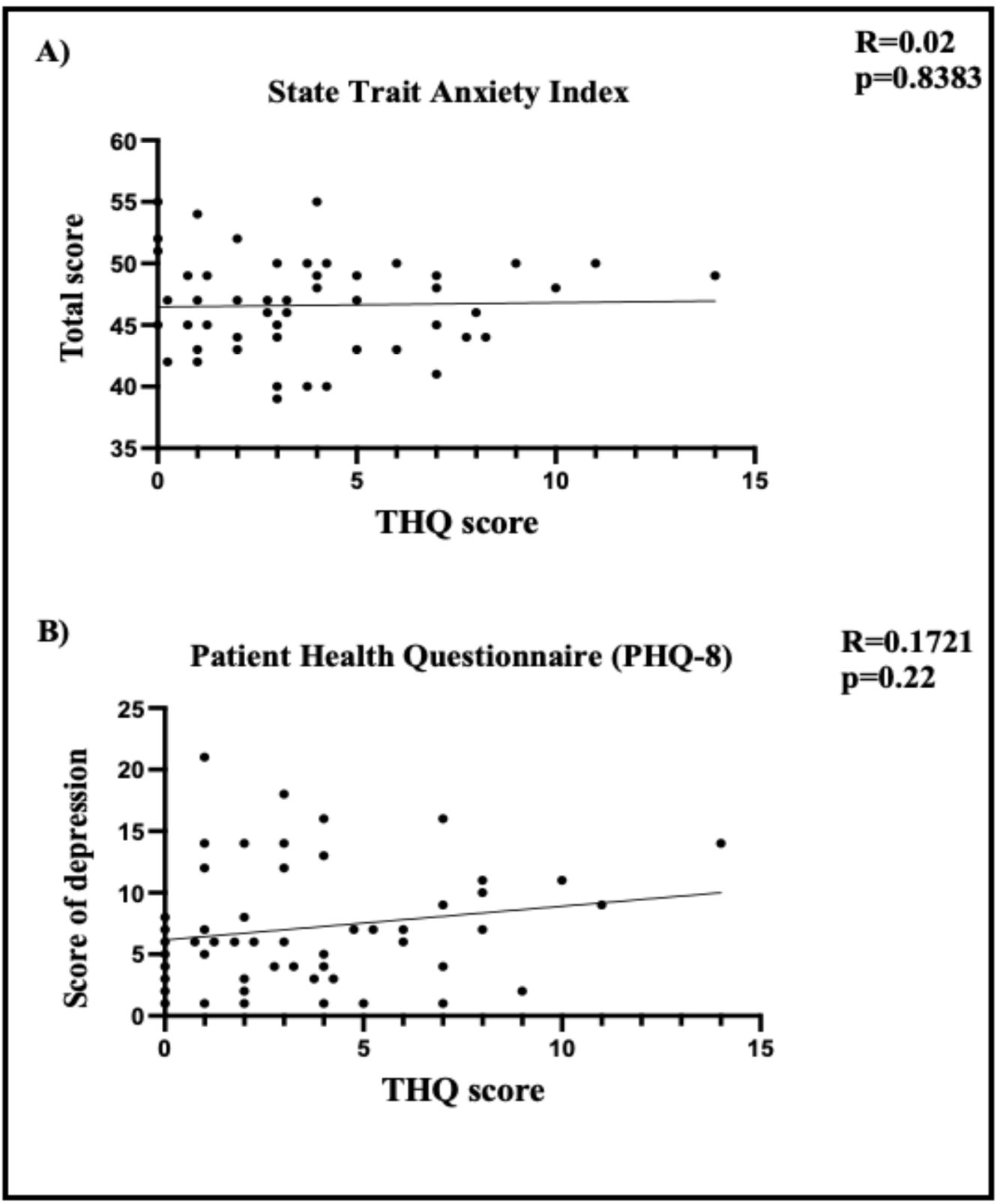
Men’s overall scores on the State Trait Anxiety Index or the Patient Health Questionnaire (PHQ-8) do not correlate with THQ score. All questionnaires of this study have been analyzed to the THQ score for each patient (State-trait anxiety index, patient health questionnaire). Trend lines represent single-variable linear regression. R= Pearson’s coefficient, with p values listed. Each point represents an individual sample. N=51

**Table 3:** State-Trait Anxiety index for the study population. For each question, each patient had to answer by one of the following answers (1. not at all, 2. somewhat, 3 moderately so and 4. very much so) from a revision of the original Form X (Spielberger et al., 1970). The answers were attributed a number score. Then, the total score for every question by THQ group was calculated as the average ± the standard deviation. The total score ranges from 20 to 80, with higher scores indicating greater anxiety. The total score for the low THQ group vs high THQ group is not significantly different (p=0.8881,Unpaired T-test, Two-tailed).

| Variable | Whole samples (n=51) | Low THQ (0-2)<br>N=19 | Medium THQ (3-7)<br>N=25 | High THQ (≥8)<br>N=7 |
| --- | --- | --- | --- | --- |
| Calm<br>Average ± SD | 3.1 ± 0.8 | 3.3 ± 0.6 | 2.9 ± 0.9 | 3 ± 0.6 |
| Secure<br>Average ± SD | 3.3 ± 0.8 | 3.4 ± 0.7 | 3.4 ± 0.9 | 3 ± 0.8 |
| Tense<br>Average ± SD | 1.9 ± 0.7 | 1.9 ± 0.6 | 2 ± 0.7 | 1.6 ± 0.8 |
| Regretful<br>Average ± SD | 1.7 ± 0.8 | 1.5 ± 0.8 | 1.7 ± 0.8 | 1.7 ± 0.8 |
| At ease<br>Average ± SD | 2.9 ± 0.8 | 3 ± 0.8 | 2.8 ± 0.8 | 3.1 ± 0.7 |
| Upset<br>Average ± SD | 1.4 ± 0.6 | 1.4 ± 0.6 | 1.4 ± 0.5 | 1.7 ± 0.8 |
| Presently worrying<br>Average ± SD | 1.9 ± 0.9 | 2.1 ± 0.9 | 1.9 ± 0.9 | 1.9 ± 1.1 |
| Rested<br>Average ± SD | 2.8 ± 0.8 | 2.9 ± 0.9 | 2.6 ± 0.8 | 2.8 ± 0.8 |
| Anxious<br>Average ± SD | 2 ± 0.7 | 2.1 ± 0.8 | 2 ± 0.6 | 1.6 ± 0.8 |
| Comfortable<br>Average ± SD | 3.1 ± 0.7 | 3.2 ± 0.7 | 2.9 ± 0.8 | 3.3 ± 0.5 |
| Self-confident<br>Average ± SD | 3.1 ± 0.8 | 3.2 ± 0.7 | 3 ± 1 | 3.3 ± 0.5 |
| Nervous<br>Average ± SD | 1.7 ± 0.7 | 1.7 ± 0.6 | 1.6 ± 0.8 | 1.7 ± 0.5 |
| Jittery<br>Average ± SD | 1.4 ± 0.7 | 1.4 ± 0.6 | 1.4 ± 0.9 | 1.4 ± 0.7 |
| High strung<br>Average ± SD | 1.7 ± 0.8 | 1.6 ± 0.8 | 1.7 ± 0.8 | 2.1 ± 1.1 |
| Relaxed<br>Average ± SD | 2.8 ± 0.8 | 2.8 ± 0.7 | 2.6 ± 0.9 | 3 ± 1 |
| Content<br>Average ± SD | 2.8 ± 0.8 | 2.9 ± 0.7 | 2.9 ± 0.8 | 2.3 ± 0.8 |
| Worried<br>Average ± SD | 2 ± 0.7 | 2 ± 0.7 | 2 ± 0.7 | 2.3 ± 1 |
| Over-exicted/rattled<br>Average ± SD | 1.3 ± 0.5 | 1.4 ± 0.7 | 1.2 ± 0.4 | 1.6 ± 0.5 |
| Joyful<br>Average ± SD | 2.9 ± 0.8 | 2.8 ± 0.8 | 3 ± 0.7 | 2.6 ± 0.8 |
| Pleasant<br>Average ± SD | 3 ± 0.7 | 3 ± 0.8 | 3 ± 0.7 | 2.9 ± 0.7 |
| Total score (20 -80)<br>Average ± SD | 46.6 ± 3.9 | 47.1 ± 4.1 | 46 ± 4 | 47.3 ± 3.9 |

**Table 4:** Patient Health questionnaire (or PHQ-8). For each question (total 8), each patient had to answer by one of the following answers (0. not at all, 1 several days, 2. more than half the time and 3. nearly every day). The answers were attributed a number score. Then, the total score for every question by THQ group was calculated as the average ± the standard deviation. For the last part, the patient needs to answer of the level of difficulty (1. not difficult, 2. somewhat, 3. moderately and 4. very much). At the end, a sum of all the points for each answer was assigned to each patient to calculate his score of depression as described in Kroenke K, Spitzer RL, Williams JB. The PHQ-9: Validity of a Brief Depression Severity Measure. J Gen Intern Med. 2001 September; 16(9): 606-613. The total score for the low THQ group vs high THQ group is not significantly different (p=0.1554, Mann–Whitney, Two-tailed).

| Variable | Whole samples (n=51) | Low THQ (0-2)<br>N=19 | Medium THQ (3-7)<br>N=25 | High THQ (≥8)<br>N=7 |
| --- | --- | --- | --- | --- |
| Q1. Little interest or pleasure in doing things<br>Average ± SD | 0.75 ± 0.81 | 0.8 ± 0.9 | 0.7 ± 0.8 | 0.9 ± 0.7 |
| Q2. Feeling down, depressed or hopeless<br>Average ± SD | 0.52 ± 0.64 | 0.45 ± 0.7 | 0.5 ± 0.7 | 0.7 ± 0.5 |
| Q3. Trouble falling or staying asleep or sleeping too much<br>Average ± SD | 0.92 ± 0.99 | 0.9 ± 1.1 | 0.8 ± 0.9 | 1.6 ± 1 |
| Q4. Feeling tired or having little energy<br>Average ± SD | 1.02 ± 0.83 | 0.95 ± 0.9 | 1.1 ± 0.8 | 0.9 ± 0.7 |
| Q5. Poor appetite or overeating<br>Average ± SD | 0.71 ± 0.94 | 0.65 ± 0.8 | 0.6 ± 0.9 | 1.1 ± 1.3 |
| Q6. Feeling bad about yourself or that you are a failure or have let yourself or your family down<br>Average ± SD | 0.63 ± 0.84 | 0.55 ± 0.7 | 0.6 ± 0.9 | 0.9 ± 1.1 |
| Q7. Trouble concentrating on things such as reading the newspaper or watching television<br>Average ± SD | 0.81 ± 0.86 | 0.6 ± 0.6 | 0.8 ± 1 | 1.3 ± 0.8 |
| Q8. Moving or speaking so slowly that other people could have noticed? Or the opposite – being so fidgety or restless that you have been moving around a lot more than usual<br>Average ± SD | 0.21 ± 0.5 | 0.1 ± 0.3 | 0.2 ± 0.5 | 0.7 ± 0.8 |
| If you checked off any problems, how difficult have these problems made it for you to do your work, take care of things at home, or get along with other people?<br>Average ± SD | 1.6 ± 0.82 | 1.7 ± 0.9 | 1.6 ± 0.7 | 1.1 ± 1.1 |
| Score of depression (0-27)<br>Minimal 0-7<br>Moderate 8-15<br>Severe 16-28<br>Average ± SD<br>N (%) | 7.17 ± 4.91<br>Minimal 33 (65)<br>Moderate 14 (27)<br>Severe 4 (8) | 6.7 ± 5.2<br>Minimal 13 (69)<br>Moderate 5 (26)<br>Severe 1 (5) | 7 ± 5<br>Minimal 18 (72)<br>Moderate 4 (16)<br>Severe 3 (12) | 9.1 ± 3.8<br>Minimal 2 (28.5)<br>Moderate 5 (71.5)<br>Severe 0 (0) |

### Anxiety-like behavior is increased in male and female offspring derived from 4 THQ-associated miRNA-injected zygotes

**Fig. 6** illustrates the effects of injecting miR-532-3p, miR-491-5p, miR-375-3p and miR-361-3p into fertilized mouse oocytes, at concentrations representative of those found in men reporting high THQ scores, on offspring behaviors compared to injecting buffer alone or a random sequence miRNA at a concentration equal to the sum of all 4 injected THQ miRNAs. Anxiety-like behavior was assessed in the elevated plus maze for both male and female mice derived from THQ miRNA injected zygotes compared to control injected ones. **Fig. 6A** shows that the percentage of open arm time (OAT%) decreased significantly in both sexes following the 4 miRNA injection, indicating increased anxiety-like behavior for both sexes. A two-way ANOVA revealed a significant main effect of the injection (treatment) on OAT (F₂,₁₂₄ = 17.57, p < 0.001), whereas neither the main effect of sex (F₁,₁₂₄ = 0.02, p = 0.89) nor the sex × treatment interaction (F₂,₁₂₄ = 1.37, p = 0.26) reached significance. Pairwise comparisons with Bonferroni correction indicated no sex differences within treatment groups; however, significant treatment effects were present in both males (F₂,₁₂₄ = 4.79, p = 0.011) and females (F₂,₁₂₄ = 9.79, p < 0.001). In males, 4 miRNA-injected animals showed lower OAT% compared with both buffer (p = 0.03) and scrambled RNA controls (p = 0.02). A similar pattern was observed in females (p = 0.042 for 4 miRNA vs. buffer; p < 0.001 for 4 miRNA vs. scrambled RNA).

**Figure. 6:**
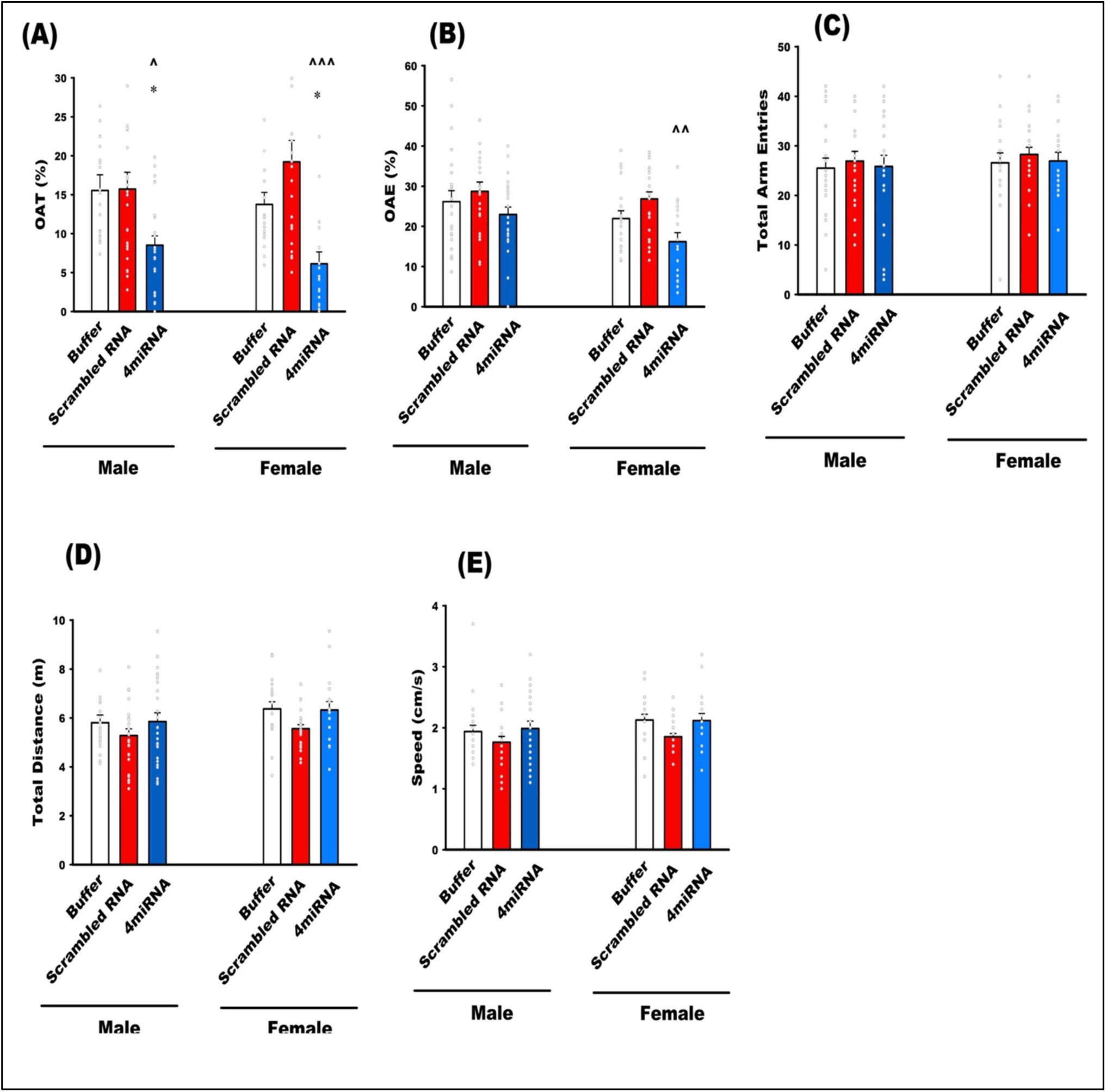
Behavioral parameters from Elevated Plus Maze (EPM) test in male and female mice derived from blastocysts injected with Buffer, Scrambled RNA, or the 4miRNA cocktail. **A**) Both males and females treated with 4miRNA showed a significant reduction in the percentage of time spent in the open arms (OAT%) compared with Buffer and Scrambled RNA controls, indicating increased anxiety-like behavior. **(B)** Females treated with 4miRNA exhibited a significant reduction in the percentage of open arm entries (OAE%) compared with Scrambled RNA controls. No significant differences were observed among treatment groups in males. **(C)** Total arm entries did not differ across treatments or sexes, indicating that locomotor initiation and general motor motivation were not affected. **(D)** Total distance traveled (m) **(E)** average speed (cm/s) did not differ among groups. Two-way ANOVA followed by Bonferroni post hoc tests was used for statistical analysis. Data are presented as mean ± SEM (Buffer: n = 22 males, 19 females; Scrambled RNA: n = 24 males, 23 females; 4miRNA: n = 25 males, 17 females) with individual animal data points overlaid. *p < 0.05 vs. Buffer; ^p < 0.05, ^^p < 0.01, ^^^p < 0.001 vs. Scrambled RNA.

The percentage of open arm entries (OAE%; **Fig. 6B**) was also altered following 4 miRNA administration, with a significant main effect of treatment (F₂,₁₂₄ = 7.04, p < 0.001) and sex (F₁,₁₂₄ = 5.65, p = 0.019). Post hoc analyses showed that this injection significantly reduced OAE% in females (F₂,₁₂₄ = 5.34, p = 0.006). Specifically, 4 miRNA-injected females exhibited lower OAE% compared with scrambled RNA controls (p = 0.004), resulting in a significant sex difference within the 4 miRNA group (F₁,₁₂₄ = 4.48, p = 0.036). In contrast, no significant effects of treatment or sex were detected in locomotor indices, including total arm entries (F₂,₁₂₄ = 0.35, p = 0.70; **Fig. 6C**), total distance traveled (F₂,₁₂₄ = 2.88, p = 0.068; **Fig. 6D**), or average speed (F₂,₁₂₄ = 2.85, p = 0.061; **Fig. 6E**).

In summary, 4 miRNA injection reduced open arm exploration (decreased OAT%) in both males and females compared with either buffer or scrambled RNA controls, consistent with increased anxiety-like behavior. A sex-dependent effect emerged in OAE%, where females, but not males, showed reduced entry frequency, suggesting a stronger avoidance strategy. Importantly, locomotor activity was unaffected, indicating that the behavioral changes reflect alterations in anxiety-like behavior rather than general activity levels.

### Depressive-like behavior is increased in male and female offspring derived from 4miRNA-injected zygotes

**Figure 7** illustrates the total immobility time (an index of depressive-like behavior) in male and female offspring subjected to the forced swimming test (FST). A two-way ANOVA revealed a significant main effect of treatment (4 miRNA, scrambled RNA, vs. buffer) on immobility time (F₂,₁₂₄ = 11.3, p < 0.001). In contrast, there was no significant main effect of sex (F₁,₁₂₄ = 0.29, p = 0.58) and no significant sex × treatment interaction (F₂,₁₂₄ = 0.03, p = 0.97). Pairwise comparisons with Bonferroni correction indicated that both male and female mice injected with 4 miRNA exhibited significantly longer immobility compared with buffer-injected or scrambled RNA controls (males: F₂,₁₂₄ = 7.09, p = 0.001; females: F₂,₁₂₄ = 4.6, p = 0.012). In males, immobility time was significantly higher in 4 miRNA-injected animals compared with both buffer (p = 0.002) and scrambled RNA controls (p = 0.013). A similar pattern was observed in females (p = 0.019 for 4miRNA vs. buffer; p = 0.036 for 4 miRNA vs. scrambled RNA).

**Figure 7.**
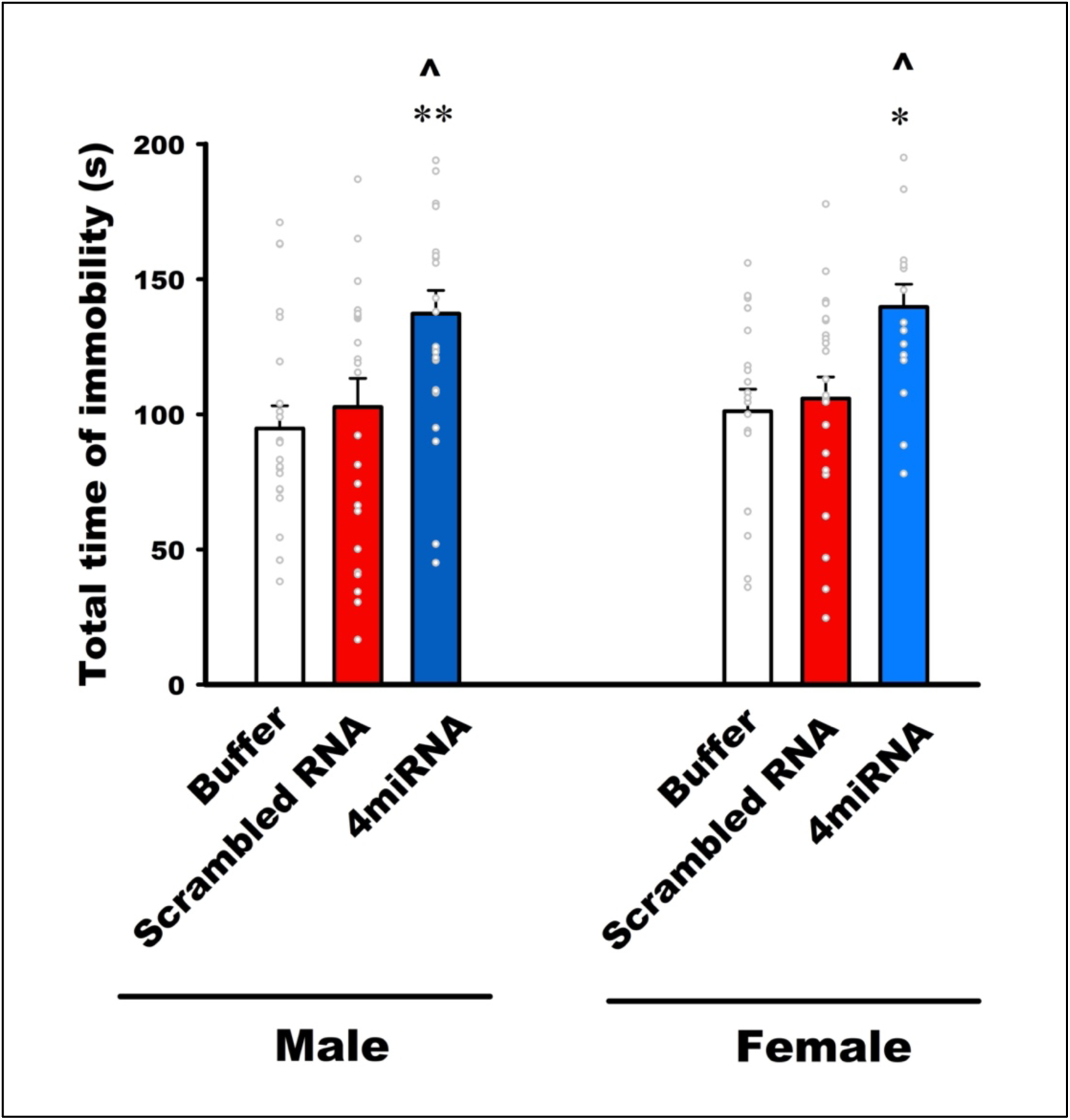
Immobility time in the forced swim test (FST) in male and female mice derived from blastocysts injected with Buffer, Scrambled RNA, or 4miRNA. The total time of immobility increased in 4miRNA group compared with both buffer and scrambled RNA control in males and females. Data are presented as mean ± SEM (Buffer: n = 22 males, 19 females; Scrambled RNA: n = 24 males, 23 females; 4miRNA: n = 25 males, 17 females) with individual animal data points overlaid. *p < 0.05, **p < 0.01 vs. Buffer; ^p < 0.05 vs. Scrambled RNA.

Together, these findings demonstrate that zygote-stage injection of a cocktail of the 4 miRNAs each at concentrations analogous to those delivered to human zygotes by men who experienced high levels of adult trauma leads to a significant increase in immobility time in the FST. When considered alongside the EPM results, the data indicates a consistent behavioral shift toward a passive stress-coping strategy and heightened vulnerability to inescapable stress (learned helplessness), features commonly associated with depressive-like phenotypes in rodent models.

## Discussion

Susceptibility to psychiatric disorders has a substantial inherited component, ranging from 50-80%, depending upon the disease (24). However, extensive genetic analysis has not fully accounted for this inheritance(25). While a component of this effect is likely a consequence of how parents raise offspring, differences in parenting alone cannot account for the observation that newborns of traumatized men already display elevated brain white matter (10). Moreover, a compelling mathematical model using summary statistics for eleven neuropsychiatric and three metabolic disorders finds that a portion of their heritability that cannot be accounted for by classical genetics may instead be attributed to epigenetic inheritance(26). But direct studies on humans across generations are inherently challenging, highlighting the importance of validating findings from model systems like chronic stress paradigms in rodents where this phenomenon has been studied extensively.

While rodent models have been instrumental in elucidating mechanisms underlying psychiatric disorders, they face inherent limitations in modeling environmentally driven conditions such as PTSD, where human traumatic experiences are heterogeneous and shaped by complex psychosocial and biological factors that cannot be fully recapitulated in animals. While rodent models of PTSD have given some insights into both the mechanisms of their effects on the traumatized rodents (27) as well as on their offspring (28), they rely on experimentally imposed stressors whose relevance to human trauma responses is necessarily approximate, and on physiological readouts that may only partially mirror those observed in humans.

This study overcomes the first challenge by focusing on the role of specific sperm miRNAs whose content is altered in men based on their responses to a PTSD-predictive questionnaire. And then we explored how these epigenetic changes affect the physiology of the next generation of mice, introducing a novel way to connect trauma responses in humans to biological processes across generations. This approach provided support for the shared property of how extreme stress alters the content of sperm miRNA in a stress-dependent manner that likely leads to distinct changes in offspring in humans, as it does in rodents.

In particular, we found that the exposure of a population of 51 men to adult trauma, like crime-related events, general disasters and traumatic experiences (e.g., injury, natural disasters, witnessing death), and unwanted physical and sexual experiences quantified by the THQ survey(29), leads to an increase in the content of a set of sperm miRNAs (elevated levels of miRs-532-3p, 491-5p, 375-3p and 361-3p) that differed from those we found altered previously in men exposed to early life trauma associated with being raised in an abusive and/or dysfunctional family quantified by the ACE survey; specifically, reduced levels of members of the miR-34/449 family (16). Interestingly, they differ from the 16 mouse sperm miRNAs implicated in transmitting across generations depression-like phenotypes in males exposed to chronic unpredictable stress (30).

As noted above, the correlation of miR-361-3p levels with THQ score fell just below the standard p-value threshold of 0.05 but the correlation coefficient was indicative of a strong association. Moreover, all of the sperm from the more than 25 men with very low THQ scores had miR-361-3p levels that were barely detectable (**Fig. 3**), whereas all 5 men that had high miR-361-3p levels in their sperm also had higher than average levels of the other 3 miRNAs, implying that elevated levels of sperm miR-361-3p likely have the potential to contribute to at least some of the stress-associated traits that might be transmitted epigenetically to offspring of these men.

miR-375-3p and miR-532-3p, are of particular interest because, among the hundreds of miRNAs found in mouse and human sperm, they are 2 of the 9 whose 3-5-fold elevated levels have been implicated by Rodgers et al. (5) in how chronic variable (CV) stress of male mice leads to the suppression of the HPA axis response in their offspring(5). Strikingly, defective HPA axis function is a common defect associated with a variety of psychiatric disorders(31, 32) including those found in offspring of parents with PTSD(33). Alternatively, suppression of the HPA axis may reflect a positive outcome, indicating the inheritance of resilience from stress. But Rodgers et al.(34) did not report individual functions for each of the 9 miRNAs. So, it is possible that miRs-375 and 532 contribute to different aspects of offspring phenotypes, particularly because their altered expression in the sperm of men with high THQ scores is associated with two other miRNAs that do not change after CV stress.

The two other miRNAs, miR-361-3p and miR-491-5p are also of particular interest. First, their rise in expression in men with high THQ scores is strong, 130X and 5X respectively. These changes are not because their elevated expression is extremely high, but because at baseline the levels of both are very low. Thus, introduction of miRNAs into zygotes upon fertilization that are not normally even present has a very high likelihood of having a major effect on embryo development. miR-361-3p has been shown to have activities that would be expected to significantly alter preimplantation embryo function when inappropriately introduced by sperm such as activating the Erk Map kinase by suppressing expression of known tumor suppressors in the context of promoting cell migration and proliferation (35). miR-491-5p also regulates cell proliferation in another setting where it enhances glioblastoma cell growth (36). Interestingly, reduced miR-361-5p has been implicated in transmission of autism (37). To determine how the elevation of these specific miRNAs affects offspring and to translate our findings, we injected these four miRNAs into control mouse zygotes at concentrations similar to what would be expected to be delivered by human sperm from men scoring high on the THQ survey. The offspring generated displayed properties consistent with elevated anxiety and depression-like behaviors in both males and females compared to control zygotes injected with either buffer alone or buffer containing a miRNA with random sequence at the same levels as the 4 THQ miRNAs combined.

The THQ survey is used to identify those with increased risk of PTSD due to trauma exposure(18), and it is known that offspring of those exposed to repeated trauma stressors are at increased risk for not only PTSD but other mental health problems (38). Yet only ∼10% of men who score high on this survey develop PTSD (39). This fact is consistent with our survey of anxiety and depression that showed most of the relatively small number of men reporting high THQ scores did not score higher than men reporting low THQ scores in either the State-Trait Anxiety Inventory or the Patient Health Questionnaire (PHQ-8). (see **Tables 3 and 4** and **Supplementary Fig. 2)**.

However, it is important to note that self-report screeners are not a substitute for clinical evaluations and may fail to capture important information about a person’s mental health picture. For instance, although there were no group differences in total PHQ-8 depression score, participants in the high THQ group scored noticeably higher on symptoms related to difficulty sleeping, appetite changes, feeling bad about oneself, difficulty concentrating, and psychomotor retardation/agitation than those in the low and medium THQ groups. Of note, these symptoms overlap with several key symptoms of PTSD (40), which, in addition to intrusive thoughts about trauma, is characterized by sleep disturbances, concentration difficulties, increased physiological arousal and reactivity, and strong negative opinions of oneself, others, and the world.

We did not administer a PTSD screener in this study, as it is unlikely that men in our sample of sperm donors would have met full diagnostic criteria for PTSD and, therefore, we anticipated that measures of anxiety and depression may be more sensitive to sub-clinical mental health experiences. It should also be stressed that the sample of the participants in the high THQ group is small, so any observations about their mental health presentation are preliminary.

Nevertheless, nearly all the men who scored high on the THQ exhibited elevated levels of all 4 sperm miRNAs. This observation, in combination with subtle changes on their clinical symptom questionnaires, supports two complementary conclusions. First, exposure of men to trauma that alters the content of specific miRNAs in their sperm can put their offspring at risk of developing psychiatric conditions, and second, there may be behavioral alterations similar to but far more subtle than outright PTSD, linking trauma exposure to the stress phenotype-inducing miRNA changes we observed. Together our findings provide a compelling avenue for future work identifying interventions that might reset these miRNA changes in highly traumatized men. Given that approximately 10% of the population we studied report THQ scores high enough induce these significant sperm miRNA changes, and a similar proportion exhibit different miRNA changes associated with high ACE scores (12), a substantial number of men may be at risk of transmitting stress-related traits to the next generation.

Future studies will evaluate whether additional phenotypes are altered on mice derived from injecting zygotes with these 4 miRNAs, as well as test the role of individual THQ associated sperm miRNAs in generating them.

## Material and Methods

### Participant recruitment and clinical assessment

All research involving human samples was approved by the Tufts University Institutional Review Board (IRB MOD-15-STUDY00001150). Written informed consent was obtained from all participants in this observational study. Demographic and health information for all samples was provided by the clinical team to the research group in a de-identified, password-protected spreadsheet to the research team, with access limited to the authors of this study. Eligible men completed the trauma history questionary (THQ) and Adverse Childhood Experience (ACE) surveys, in a self-reporting format following informed consent. For the THQ, each “yes” response was assigned one point with multiple encounters contributing additional points. In addition, each participant completed the state-trait anxiety inventory, the Patient Health Questionnaire (PHQ-8) and a resilience questionnaire. Men then provided a semen sample as part of their initial workup (n = 51). All participants had normal sperm count and motility as defined by WHO parameters (Table 1). Mature motile sperm were isolated using a Percoll gradient to effectively separate mature sperm from immune cells, dead tissues, immature sperm, and other contaminants (41). All participants were compensated with a $50 gift card. Recruitment and sample collection were supported by funding allocated for a three-year enrollment period

### RNA extraction and quantitative real-time PCR

Total RNA was extracted from human sperm samples using the miRVana miRNA Isolation kit (Invitrogen) according to the manufacturer’s protocol. RNA concentration and quality were assessed using NanoDrop-1000 (Thermofisher Scientific). An aliquot of 10ng of total RNA from each sperm sample was used for cDNA synthesis with the TaqMan Advanced cDNA synthesis kit (Applied Biosystems).). Real-time PCR was performed in duplicate using a StepOnePlus Real-Time PCR system with TaqMan Master Mix (Applied Biosystems). Relative expression levels were calculated using the comparative ΔΔCt method, with miR-192-5p serving as the internal control for all samples. The miRNA primers used in this study were TaqMan^TM^ Advanced miRNA assay (#A25576, Thermofisher, miR-34c-5p, #478052_miR, miR-192-5p, #478262_miR, miR-532-3p, #478336_miR, miR-375-3p, #478074_miR, miR-361-3p, #478055_miR, miR-491-5p, miR-478132_miR, miR-193b-3p, #478314_miR, miR-99a-5p, #478519_miR, miR-7-5p, #483061_miR, miR-147b-5p, #483120_miR).

miRNA Sequencing Data Processing and Differential Expression Analysis After RNA extraction, samples from men with low THQ score (n=5) and high THQ score (n=5) were selected for miRNA sequencing. All extracted RNA was used for library preparation following Illumina’s TruSeq-small-RNA-sample preparation protocols (Illumina, San Diego, CA, USA). Library quality control and quantification were performed using Agilent Technologies 2100 Bioanalyzer High Sensitivity DNA Chip. Single end 50bp sequencing was carried out on the Illumina’s Hiseq 2500 platform following the manufacture’s recommended protocols. Raw sequencing reads were processed using the in-house ACGT101-miR pipeline (LC Sciences, Houston, Texas, USA) to remove adapter dimers, low-quality reads, low-complexity sequences, and other RNA species (including rRNA, tRNA, snRNA, and snoRNA) and repetitive elements. Clean reads with length between 18 and 26 nucleotides were retained for downstream analysis. Unique sequences were aligned to species specific precursors miRNA described in miRBase version 22.0 using BLAST to identify known miRNAs as well as novel 3p- and 5p-derived miRNAs. During alignment, length variation at both 3’ and 5’ ends and up to one internal mismatch were permitted. Sequences mapping to annotated mature miRNAs located on the hairpin arms of species-specific precursors were classified as known miRNAs. Sequences mapping to the opposite arm of a known precursor hairpin were considered candidate novel 5p- or 3p-derived miRNA. Remaining sequences were subsequently aligned to precursor miRNAs from other species in miRBase 22.0 (excluding the target species), and the resulting mapped precursors were further aligned to the specific species genomes to determine their genomic locations. These miRNAs were also classified as known miRNA. Unmapped sequences were aligned to the reference genomes, and the potential precursor hairpin structures were predicted from 80-nucleotide flanking sequences using RNAfold software (http://rna.tbi.univie.ac.at/cgi-bin/RNAfold.cgi). Secondary, structure prediction was evaluated based on the following criteria: (1) ≤12 per bulge in the stem region) (2) ≥16 base pairs in the stem region) (3) minimum free energy ≤-15 kcal/mol (4) total hairpin length (including stem and terminal loop) ≥50 (5) hairpin loop length ≤20. (6) ≤8 nucleotides per bulge in mature region) (7) ≤4 biased errors per bulge in the mature region (8) ≤2 biased bulges in the mature region (9) ≤7 total mismatches in the mature region; (10) ≥12 base pairs in the mature region and (11) ≥80% of the mature miRNA sequence located within the stem regions.

### Microinjection

All experimental protocols were approved by the MaineHealth Institute for Research Institutional Animal Care and Use Committee and conducted in an AAALAC accredited facility in accordance with animal care guidelines. Mice were maintained at a 12-hour (6AM-6PM) light-dark cycle and fed a rodent diet (Research Diets, Teklad 2918 Chow Diet) *ad libitum* in a pathogen-free room.

Overall, we followed the procedure used by Radger et al (5) who replicated the transmission of altered HPA axis response of mice exposed to chronic variable stress by injecting the 9 sperm miRNAs (0.11 ng each in a total of 1ul) they observed elevated in their sperm into fertilized zygotes of control mice.

Thus, we used the same control mice, C57BL/6J 3-week-old females (approximately 12 grams), superovulated via intraperitoneally injections of pregnant mare serum (5 IU, ProspecBio) followed by human chorionic gonadotropin (5 IU, ILEX Bioscience). The superovulated donor females were then mated with 129S6/SvEvTac stud males to produce C57BL/6J:129S6/SvEvTac hybrid mouse zygotes. These F1 hybrid fertilized zygotes were collected 22-24 hours after HCG treatment and randomly assigned for cytoplasmic microinjection of the three conditions, buffer alone, a single miRNA control, and a pool of our 4 miRNAs. Microinjection into the cytoplasm of zygotes was performed using Zeiss Axiovert S100 inverted microscope with Eppendorf TransferMan NK controllers and an Eppendorf Femtojet. Each injection was deemed successful after visual swelling in the cytoplasm and survival of the injected zygote.

The experimental condition we used that attempted to reflect what occurs in men with high THQ scores considered not only the fold-change observed for each miRNA shown in **Fig. 3** but also the large differences in their baseline levels (miR-491-5p and miR-361-3p being extremely low compared to the others and miR-532 being much higher than miR-375). Specifically, mouse miR-532 and miR-375, both of which were among the 9 miRNAs studied by Rodgers, were injected at twice (**0.2 ng/μL**) and seven times (**0.7 ng/μL**), respectively, the amounts they used to account for our finding larger differences for each. Unlike any of the 9 miRNAs they used, and we found that miR-361 was barely detectable and miR-491was so low as to not be easily quantified in mouse sperm, and we find it barely detectable in human sperm. Thus, we injected **0.05 ng/μL** of both to approximate the strong increase we detected in human sperm from men with high THQ scores. Together the miRNAs were 1ng/μL total concentration in MIJ Buffer (1 mM Tris-HCl (pH 7.5), 0.1 mM EDTA). The control conditions included a scrambled miRNA to reflect the cumulative increase of nonspecific miRNAs in the zygote, and MIJ buffer alone, used to control for potential effects of zygote disturbance caused by the injection procedure.

Following the microinjection procedure, the surviving injected zygotes were implanted into 0.5 day recipient females but if set up did not allow, zygotes were cultured overnight in potassium-supplemented simplex optimized media and transferred as 2-cells in KSOM, EMD media (Millipore). To prepare recipients, Swiss Webster females of reproductive age were mated to vasectomized Swiss Webster males and identified the following day as a recipient via a copulation plug. Each female was implanted with between 14 and 30 zygotes of a single condition group. Recipients of the same condition were cohoused or if only one recipient per condition they were single housed during pregnancy. Pups were weaned at post-natal day 28 into cages with the same sex (up to four mice per cage).

### Behavioral Tests

To assess behavioral changes, the Elevated Plus Maze (EPM) and Forced Swim Test (FST) were performed on animals at least 8 weeks of age. Animals were transferred to the behavioral testing room one hour prior to testing for habituation. All experiments were conducted between 8:00 a.m. and 4:00 p.m.

### Elevated Plus Maze (EPM)

The EPM is a well-established test for assessing anxiety-like behavior in rodents, based on their natural aversion to open spaces and preference for enclosed areas(42). The apparatus consisted of a plus-shaped maze elevated 40 cm above the floor, with two open arms and two closed arms. Mice were placed in the center of the maze and allowed to explore for 5 minutes. Behavioral parameters, including time spent in each arm, number of entries into each arm, total distance traveled, and mean speed, were recorded using ANY-maze software (Stoelting Co., USA). For analysis, the following indices were calculated: percentage of open arm time (OAT% = [time in open arms / total time in open + closed arms] × 100), percentage of closed arm time (CAT% = [time in closed arms / total time in open + closed arms] × 100), and percentage of open arm entries (OAE% = [entries into open arms / total arm entries] × 100), along with locomotor activity measures (total arm entries, speed, and total distance).

### Forced Swim Test (FST)

The FST is a widely used paradigm to evaluate depressive-like behavior in rodents(43). The apparatus consisted of a Plexiglas cylinder (20 cm diameter, 30 cm height) filled with water to a depth of 15 cm, maintained at 24 ± 1 °C. Each mouse was placed individually in the cylinder for 6 minutes. Immobility was recorded during the final 4 minutes of the test, defined as the absence of active movements except those necessary to keep the head above water. Immobility duration (in seconds) was automatically quantified using ANY-maze software.

## Statistical analysis

Demographic, psychological, and sperm characteristics were summarized by THQ group (low: 0-2, medium: 3-7, high: ≥8) using appropriate summary statistics for normally and non-normally distributed data. For miRNA expression analyses, differential expressions from sequencing data were evaluated using **Fisher’s exact test** in the LC Sciences pipeline, with p-values adjusted for multiple comparisons using the **Benjamini-Hochberg false discovery rate (FDR)**; adjusted p < 0.05 was considered significant. For qPCR, comparisons between low (0–2) and high (≥8) THQ groups were performed using **unpaired t-tests** for normally distributed miRNAs (miR-532-3p, miR-491-5p) and **Mann-Whitney tests** for non-normal miRNAs (miR-361-3p, miR-375-3p). Correlation analyses were performed using reverse ΔCt values from qPCR. Associations between miRNA expression and THQ or ACE scores were evaluated using Spearman’s correlation, with linear regression lines fitted for visualization. Pearson’s correlation was used for the State-Trait Anxiety Inventory and PHQ-8 scores. Animal behavioral data was analyzed using two-way ANOVA followed by Bonferroni-corrected post-hoc tests. All analyses were conducted in GraphPad Prism v10, with p < 0.05 considered statistically significant.

## Acknowledgments.

We acknowledge Dr. Emily Feig, Dept of Psychiatry, Mass General Hospital, Boston MA, for recommending using the Anxiety and Depression Questionnaires in this study.

We acknowledge the Mouse Genome Modification Core Facility at the MaineHealth Institute for Research for their expert support in providing in vitro fertilization, embryo transplantation, and incubation to the blastocyst stage for in vitro experiments. We are grateful to Logan Douglas at MaineHealth Institute for Research, for facilitating the performance of the behavioral tests.

## REFERENCES

1. Fitz-James MH, Cavalli G. Molecular mechanisms of transgenerational epigenetic inheritance. Nat Rev Genet. 2022;23(6):325–41.

2. Yang CH, Rando OJ. Paternal Effects in Mammals: Challenges and Opportunities. Annu Rev Biochem. 2025;94(1):253–78.

3. Saavedra-Rodriguez L, Feig LA. Chronic social instability induces anxiety and defective social interactions across generations. Biol Psychiatry. 2013;73(1):44–53.

4. Champroux A, Tang Y, Dickson DA, Meng A, Harrington A, Liaw L, et al. Transmission of reduced levels of miR-34/449 from sperm to preimplantation embryos is a key step in the transgenerational epigenetic inheritance of the effects of paternal chronic social instability stress. Epigenetics. 2024;19(1):2346694.

5. Rodgers AB, Morgan CP, Leu NA, Bale TL. Transgenerational epigenetic programming via sperm microRNA recapitulates effects of paternal stress. Proc Natl Acad Sci U S A. 2015;112(44):13699–704.

6. Tolkunova K, Usoltsev D, Moguchaia E, Boyarinova M, Kolesova E, Erina A, et al. Transgenerational and intergenerational effects of early childhood famine exposure in the cohort of offspring of Leningrad Siege survivors. Sci Rep. 2023;13(1):11188.

7. Dashorst P, Mooren TM, Kleber RJ, de Jong PJ, Huntjens RJC. Intergenerational consequences of the Holocaust on offspring mental health: a systematic review of associated factors and mechanisms. Eur J Psychotraumatol. 2019;10(1):1654065.

8. Amstadter AB, Abrahamsson L, Cusack S, Sundquist J, Sundquist K, Kendler KS. Extended Swedish Adoption Study of Adverse Stress Responses and Posttraumatic Stress Disorder. JAMA Psychiatry. 2024;81(8):817–24.

9. Chou PC, Huang YC, Yu S. Mechanisms of Epigenetic Inheritance in Post-Traumatic Stress Disorder. Life (Basel). 2024;14(1).

10. Karlsson H, Merisaari H, Karlsson L, Scheinin NM, Parkkola R, Saunavaara J, et al. Association of Cumulative Paternal Early Life Stress With White Matter Maturation in Newborns. JAMA Netw Open. 2020;3(11):e2024832.

11. Brikell I, Ruck C, Kuja-Halkola R. Genes and Rearing-What Explains Intergenerational Transmission of PTSD? JAMA Psychiatry. 2024;81(8):747–8.

12. Marcho C, Oluwayiose OA, Pilsner JR. The preconception environment and sperm epigenetics. Andrology. 2020;8(4):924–42.

13. Morgan CP, Shetty AC, Chan JC, Berger DS, Ament SA, Epperson CN, et al. Repeated sampling facilitates within- and between-subject modeling of the human sperm transcriptome to identify dynamic and stress-responsive sncRNAs. Sci Rep. 2020;10(1):17498.

14. Anda RF, Brown DW, Felitti VJ, Bremner JD, Dube SR, Giles WH. Adverse childhood experiences and prescribed psychotropic medications in adults. Am J Prev Med. 2007;32(5):389–94.

15. Tuulari JJ, Bourgery M, Iversen J, Koefoed TG, Ahonen A, Ahmedani A, et al. Exposure to childhood maltreatment is associated with specific epigenetic patterns in sperm. Mol Psychiatry. 2025.

16. Dickson DA, Paulus JK, Mensah V, Lem J, Saavedra-Rodriguez L, Gentry A, et al. Reduced levels of miRNAs 449 and 34 in sperm of mice and men exposed to early life stress. Transl Psychiatry. 2018;8(1):101.

17. Champroux A, Sadat-Shirazi M, Chen X, Hacker J, Yang Y, Feig LA. Astrocyte-derived exosomes regulate sperm miR-34c levels to mediate the transgenerational effects of paternal chronic social instability stress. Epigenetics. 2025;20(1):2457176.

18. Hooper LM, Stockton P, Krupnick JL, Green BL. Development, Use, and Psychometric Properties of the Trauma History Questionnaire. J Loss Trauma. 2011;16(3):258–83.

19. Dube SR, Anda RF, Felitti VJ, Chapman DP, Williamson DF, Giles WH. Childhood abuse, household dysfunction, and the risk of attempted suicide throughout the life span: findings from the Adverse Childhood Experiences Study. JAMA. 2001;286(24):3089–96.

20. Golier JA, Yehuda R, Bierer LM, Mitropoulou V, New AS, Schmeidler J, et al. The relationship of borderline personality disorder to posttraumatic stress disorder and traumatic events. Am J Psychiatry. 2003;160(11):2018–24.

21. Najavits LM, Weiss RD, Shaw SR, Muenz LR. “Seeking safety”: outcome of a new cognitive-behavioral psychotherapy for women with posttraumatic stress disorder and substance dependence. J Trauma Stress. 1998;11(3):437–56.

22. Knowles KA, Olatunji BO. Specificity of trait anxiety in anxiety and depression: Meta-analysis of the State-Trait Anxiety Inventory. Clin Psychol Rev. 2020;82:101928.

23. Levis B, Fischer F, Benedetti A, Thombs BD. PHQ-8 scores and estimation of depression prevalence. Lancet Public Health. 2021;6(11):e793.

24. Pettersson E, Lichtenstein P, Larsson H, Song J, Attention Deficit/Hyperactivity Disorder Working Group of the iPsych-Broad-Pgc Consortium ASDWGoti-B-PGCCBDWG, Tourette Syndrome Working Group of the Pgc SCSUDWGotPGC, et al. Genetic influences on eight psychiatric disorders based on family data of 4 408 646 full and half-siblings, and genetic data of 333 748 cases and controls - CORRIGENDUM. Psychol Med. 2019;49(2):351.

25. Grezenko H, Ekhator C, Nwabugwu NU, Ganga H, Affaf M, Abdelaziz AM, et al. Epigenetics in Neurological and Psychiatric Disorders: A Comprehensive Review of Current Understanding and Future Perspectives. Cureus. 2023;15(8):e43960.

26. Monaco AP. An epigenetic, transgenerational model of increased mental health disorders in children, adolescents and young adults. Eur J Hum Genet. 2021;29(3):387–95.

27. Aspesi D, Pinna G. Animal models of post-traumatic stress disorder and novel treatment targets. Behav Pharmacol. 2019;30(2 and 3-Spec Issue):130–50.

28. Yamakawa GR, Freeman J, Harris S, Sgro M, Vlassopoulos E, Li CN, et al. Intergenerational influences of paternal combat-related trauma on offspring behavioral and brain function. Neurobiol Stress. 2025;39:100767.

29. Bengtson L, Aubuchon-Endsley N, Meotti S, Lynch S. Trauma History Questionnaire: validation with novel samples of incarcerated women and perinatal women. Women Health. 2024;64(5):380–91.

30. Wang Y, Chen ZP, Hu H, Lei J, Zhou Z, Yao B, et al. Sperm microRNAs confer depression susceptibility to offspring. Sci Adv. 2021;7(7).

31. Juruena MF, Bocharova M, Agustini B, Young AH. Atypical depression and non-atypical depression: Is HPA axis function a biomarker? A systematic review. J Affect Disord. 2018;233:45–67.

32. Altamura AC, Boin F, Maes M. HPA axis and cytokines dysregulation in schizophrenia: potential implications for the antipsychotic treatment. Eur Neuropsychopharmacol. 1999;10(1):1–4.

33. Bowers ME, Yehuda R. Intergenerational Transmission of Stress in Humans. Neuropsychopharmacology. 2016;41(1):232–44.

34. Rodgers AB, Morgan CP, Bronson SL, Revello S, Bale TL. Paternal stress exposure alters sperm microRNA content and reprograms offspring HPA stress axis regulation. J Neurosci. 2013;33(21):9003–12.

35. Li Y, Fan L, Yan A, Ren X, Zhao Y, Hua B. Exosomal miR-361-3p promotes the viability of breast cancer cells by targeting ETV7 and BATF2 to upregulate the PAI-1/ERK pathway. J Transl Med. 2024;22(1):112.

36. Jie XF, Li YP, Liu S, Fu Y, Xiong YY. miR-491-5p regulates the susceptibility of glioblastoma to ferroptosis through TP53. Biochem Biophys Res Commun. 2023;671:309–17.

37. Yilmaz Sukranli Z, Korkmaz Bayram K, Mehmetbeyoglu E, Doganyigit Z, Beyaz F, Sener EF, et al. Trans Species RNA Activity: Sperm RNA of the Father of an Autistic Child Programs Glial Cells and Behavioral Disorders in Mice. Biomolecules. 2024;14(2).

38. Seng JS, D’Andrea W, Ford JD. Complex Mental Health Sequelae of Psychological Trauma Among Women in Prenatal Care. Psychol Trauma. 2014;6(1):41–9.

39. Charlson F, van Ommeren M, Flaxman A, Cornett J, Whiteford H, Saxena S. New WHO prevalence estimates of mental disorders in conflict settings: a systematic review and meta-analysis. Lancet. 2019;394(10194):240–8.

40. American Psychiatric Association. Diagnostic and statistical manual of mental disorders: DSM-5-TR. Fifth edition, text revision. ed. Washington, DC: American Psychiatric Association Publishing; 2022. lxix, 1050 pages p.

41. Moohan JM, Lindsay KS. Spermatozoa selected by a discontinuous Percoll density gradient exhibit better motion characteristics, more hyperactivation, and longer survival than direct swim-up. Fertil Steril. 1995;64(1):160–5.

42. Bourin M, Petit-Demoulière B, Nic Dhonnchadha B, Hascöet M. Animal models of anxiety in mice. Fundamental & clinical pharmacology. 2007;21(6):567–74.

43. Yankelevitch-Yahav R, Franko M, Huly A, Doron R. The forced swim test as a model of depressive-like behavior. J Vis Exp. 2015(97).

