## Supplemental Table 1 for "Exposure of Men to PTSD-Promoting Trauma Elevates Levels of Sperm miRNAs with Anxiety and Depression-Inducing Activities"

**Supplementary Table**

| Variable | Whole sample (n=51) | Low THQ (0-2) N=19 | Medium THQ (3-7) N=25 | High THQ (≥8) N=7 | High/low THQ group difference Mann Whitney U test |
|---|---|---|---|---|---|
| **Race/ethnicity % (n)** | **White 75 (38)<br>Asian 16 (8)<br>Black 2 (4)<br>Other 6 (3)** | **White 63 (13)<br>Asian 32 (6)<br>Black 5 (1)** | **White 80 (20)<br>Asian 4 (1)<br>Black 4 (1)<br>Other 12 (3)** | **White 86 (6)<br>Asian 14 (1)** | / |
| **Age (years), average ±SD** | **36.4 ± 2.8** | **32.6 ± 3.6** | **34;2 ± 3.9** | **36.4 ± 3.9** | **P=0.0182** |
| **BMI (kg/m²), average ±SD** | **29.6 ± 8** | **28.9 ± 6.7** | **29.9 ± 9.9** | **30.6 ± 12** | **P=0.5858** |
| **Ever smoked, n (%)** | **13 (25)** | **2 (11)** | **8 (40)** | **3 (43)** | / |
| **Current smoker, n (%)** | **9 (18)** | **2 (11)** | **5 (20)** | **2 (29)** | / |
| **Alcoholic drinks/wk median (range)** | **7 (8)** | **4 (8)** | **7 (6)** | **5 (2)** | / |
| **Illegal drug use, n (%)** | **9 (18)** | **3 (15)** | **5 (20)** | **1 (14)** | / |
| **Variable** | **Whole sample (n=45)** | **Low THQ (0-2) N=17** | **Medium THQ (3-7) N=22** | **High THQ (≥8) N=6** | **High/low THQ group difference Unpaired T-Test** |
| **Sperm count (mil/ml), mean ± SD** | **74.8 ± 51.8** | **70.8 ± 52.6** | **86.8 ± 52.5** | **42.5 ± 35.2** | **P=0.235** |
| **Sperm motility (% motile), median (range)** | **60.5 (12-91)** | **58.5 (12-80)** | **63 (32-91)** | **60.5 (35-83)** | / |
| **Sperm morphology (% normal), median (range)** | **2.5 (0-7)** | **2.5 (0-6)** | **2 (0-7)** | **2.5 (0-4)** | / |
| **Rounds cell (%) median (range)** | **0 (0-7)** | **0 (0-6)** | **1 (0-7)** | **0 (0-3)** | / |
| **Head defects (%), median (range)** | **97 (83-100)** | **97 (87-100)** | **97 (83-100)** | **92 (83-100)** | / |
| **Tail defects (%), median (range)** | **5 (0-16)** | **1 (0-9)** | **1 (0-13)** | **5 (0-16)** | / |
| **Viscosity (unitless)** | **None 29<br>Slight 9<br>Moderate 5<br>Extreme 2** | **None 12<br>Slight 1<br>Moedrate 3<br>Extreme 1** | **None 13<br>Slight 6<br>Moderate 2<br>Extreme 1** | **None 4<br>Slight 2** | / |

**Supplemental Table 1**: **Demographic, behavioral and sperm characteristic of the study population, stratified by THQ score.** Compiled data for all the 51 human samples used in this study.
